# Non-invasive measurement of metabolic rates of free-swimming jellyfish using volumetric laser scanning

**DOI:** 10.1101/2025.06.27.662033

**Authors:** Simon R. Anuszczyk, John O. Dabiri

## Abstract

Measuring energy consumption of marine organisms often requires enclosing the animal in a sealed chamber of similar volume to quantify changes in oxygen concentration of the surrounding water. This method limits measurements of free-swimming organisms without boundary effects or movement restrictions. We present energy consumption measurements of free-swimming jellyfish based on 3D morphological reconstructions of the animal during the process of catabolysis utilizing a non-invasive technique. We used a 6-meter tall, 13,600 liter water tank for swimming experiments without food present in the water column. To study continuous swimming, we implemented an onboard microelectronic swim controller that externally stimulated jellyfish swimming for 50 hours. Computer vision-based feedback control was used to enable continuous swimming against a flow current without encountering the vertical limits of the tank over a distance of 2.55 km or 15,000 body lengths. Using a 3D laser scanning system, we created morphological reconstructions of the jellyfish before, during, and after swimming experiments. Changes in animal volume were converted to energy consumption using the body chemical composition determined with elemental analysis. Free-swimming, electrically stimulated animals were found to consume 2.5 times more energy than similarly stimulated animals in a constrained environment, likely due in part to additional hydrodynamic drag associated with free-swimming. Simplified drag models could not fully account for the increased energy consumption. This highlights the need for hydrodynamic models that more effectively capture these dynamics. These measurements suggest that this technique could be applied more broadly to measuring free-swimming metabolic rates of other gelatinous invertebrates.

**SUMMARY STATEMENT:** This article presents measurements of the energy consumption of free-swimming jellyfish by converting changes in animal volume to metabolic rates using a non-invasive laser-scanning technique.

## INTRODUCTION

Energy sourcing, consumption, and allocation are vital to maintaining life. Measuring energy consumption enables the prediction of an organism’s performance and contributes to the characterization of survival, fitness, and phenology (de Groot et al., 2024). Thus, accurate energy consumption measurements are important not only for understanding physiology, but organismal biology more broadly. A variety of techniques have been developed to measure the metabolism of marine organisms. The most common of these techniques, respirometry, measures the respiratory rate, which is a metabolic measurement of oxygen consumption commonly used as a proxy for energy consumption. Respirometry comprises three main techniques each with associated benefits and drawbacks (Svendsen et al., 2015). Closed system respirometry encloses the animal in a tank, generally of a similar size as the animal, and measures the decreasing oxygen concentration in the water as the animal respires (Keys, 1930). These systems are simple and ubiquitous, but avoiding hypoxic effects and the buildup of organic waste in the chamber limits experimental duration. Flow-through respirometry instead provides a continuous flow of water through the chamber and measures the difference in oxygen concentration at the inlet and outlet to calculate oxygen consumption. While this technique minimizes hypoxic effects, concerns of sensor issues, mixing, and waste buildup due to imperfect mixing persist. Finally, intermittent-flow respirometry combines periods of closed system respirometry with flushes of the chamber to eliminate both hypoxic effects and the buildup of waste (Svendsen et al., 2015). However, all of these methods rely on changes in oxygen consumption measurable above background noise. In practice, each requires comparatively small experimental chambers which introduce other confounding variables such as movement restriction and accompanying animal stress (Ikeda et al., 2000).

Other techniques attempt to avoid the limitations of small experimental chambers by studying animal energy consumption in situ. Doubly labeled water is the standard technique for terrestrial organisms and involves introducing hydrogen and oxygen isotopes to the animal whose elimination rates are then measured. However, this technique is not applicable to marine organisms due to high body water turnover rates making elimination rate measurements infeasible (Treberg et al., 2016). Biotelemetry, including electromyogram sensors and accelerometry, has been used to indirectly estimate energy consumption. These techniques are most applicable to large fish due to tag size and generally do not control for different types of activity. Like doubly labeled water, isotopic trace turnover methods such as carbon isotopes consumed through feeding are valuable for energy consumption measurements. However, these techniques require extensive validation before application to measure decay curves for each species (Treberg et al., 2016). Vertebrates have calcium carbonate structures known as otoliths as part of the vestibular system. As the otolith grows with the animal, mineral layers are deposited in accordance with both environmental conditions and metabolic conditions. In bony fishes, the otolith-isotope method can identify metabolic rates through animal dissection and analysis. This method usually involves euthanizing the animal, limiting ongoing studies; gelatinous zooplankton have no similar structures for analysis (de Groot et al., 2024). Although these techniques enable energetic measurements in more realistic free-swimming environments, their accompanying limitations make them unsuitable for gelatinous zooplankton.

Here, we develop, characterize, and implement a non-invasive technique that overcomes these limitations by inferring the energy consumption of free-swimming *Aurelia aurita* jellyfish medusae based on changes in body volume due to catabolysis. Catabolysis is the process by which an organism breaks down tissue when deprived of nutrients. Decreases in size during periods of catabolysis have been documented in many marine organisms, including fishes (Gingerich et al., 2010; Einen et al., 1998; Simpkins et al., 2003), flatworms (Gambino et al., 2023), and jellyfish (Lilley et al., 2014; Arai 1986; Ishii and Bamstedt, 1998; Fu et al., 2014). Previous work has shown starving *Aurelia aurita* jellyfish mass decreases by up to 13.4% per day (Ishii and Bamstedt, 1998) and some work predicts a derived respiratory carbon demand of more than 17% per day (Bondyale-Juez et al., 2022). One study inferred metabolic rates from average mass changes of *Aurelia aurita* ephyrae to estimate the respirometry rate. However, this study relied on euthanasia as part of their protocol for measuring animal mass limiting applicability of this technique (Fu et al., 2014). Recent developments in volumetric (3D) laser scanning enable detailed 3D reconstructions of gelatinous zooplankton (Fu et al., 2021; Daniels et al., 2023). This allows for measurements of changing body size due to catabolysis while in situ, which can be converted to energetic consumption rates (Ikeda et al., 2000).

One important confounding variable in energetics is the activity level of the animal. Jellyfish pulse rate, and thus metabolism, varies significantly based on environmental conditions and animal morphology (Tills et al., 2016; Dillon 1977; McHenry and Jed, 2003). In order to control this variable, we utilized onboard electrical stimulation (Xu and Dabiri, 2020a) to fix the animal pulse rate at a defined frequency across animals and throughout the duration of the test.

In this paper, we first describe the calibration and validation of the volumetric measurements obtained from the 3D laser scanning technique with wet weight measurements. Volumetric changes due to catabolysis detected by the laser scanning system were related to respiratory rate using established relationships in the literature and verified against closed-system respirometry measurements for both naturally swimming and electrically stimulated swimming. We conducted body composition measurements to inform this relationship. This technique was then utilized for free-swimming measurements of electrically stimulated jellyfish. Testing over swimming durations greater than 15,000 body lengths enabled assessment of long-term changes in metabolism during catabolysis. We compared these measurements against predictions from a physics-based energetics model and basal metabolism experiments.

## MATERIALS AND METHODS

### Animal hubandry

*Aurelia aurita* medusae were obtained from Cabrillo Marine Aquarium (San Pedro CA, USA) and Aquarium of the Pacific (Long Beach, CA). They were housed in a 453 liter psuedokreisel tank (Jelliquarium 360, Midwater Systems, Thousand Oaks, CA, USA). The tank was filled with artificial seawater mixed with sea salt (Instant Ocean Sea Salt, Spectrum Brands, Blacksburg, VA, USA) and deionized water at 35 parts per thousand (PPT) and kept at 21°C. The jellyfish were hand-fed twice daily with live *Artemia franciscana* brine shrimp (Hatching Shell-Free Brine Shrimp Eggs E-Z Egg, Brine Shrimp Direct, Ogden, UT, USA). Brine shrimp were hatched every other day in aerated hatchery cones of artificial seawater heated to 28°C for 24 hours, then transferred to beakers at 21°C. Brine shrimp were fed PHYTO-Feast (Reef Nutrition, Reed Mariculture Inc. Campbell, CA) and enriched with SELCO (Self-Emulsifying Lipid Concentrate, Brine Shrimp Direct, Ogden, UT, USA).

### Laser scanning for volumetric reconstruction

Volumetric reconstructions of the jellyfish were created using a single-camera laser scanning system (Fig. 1A). This system used a 671 nm continuous laser (5 W Laserglow LRS0671 DPSS Laser System) and a series of optics to generate a scanning laser sheet. The optics consisted of a mirror mounted on a rotating galvanometer (Thorlabs GVS211/M, 10 kHz bandwidth) controlled with an arbitrary function generator (Tektronix AFG3011C) to sweep across the volume of inquiry at 10 Hz; a 25 cm diameter condenser lens to redirect the laser normal to the scanning direction; and a cylindrical, 16-mm diameter glass rod used to form the laser beam into a thin sheet. The laser sheet was set to sweep normal to its plane across a distance of 11.7 cm. A high-speed camera recording at 6000 fps (Photron FASTCAM SA-Z) thus captured 600 1024 × 850 pixel or 22.5 × 18.7 cm 2D images for each scan. A custom MATLAB code was written to reconstruct the jellyfish from these image stacks. This code identified the jellyfish and performed binarization, thresholding, and morphological opening and closing operations to create a detailed reconstruction (Fig. 1B).

**Fig. 1.**
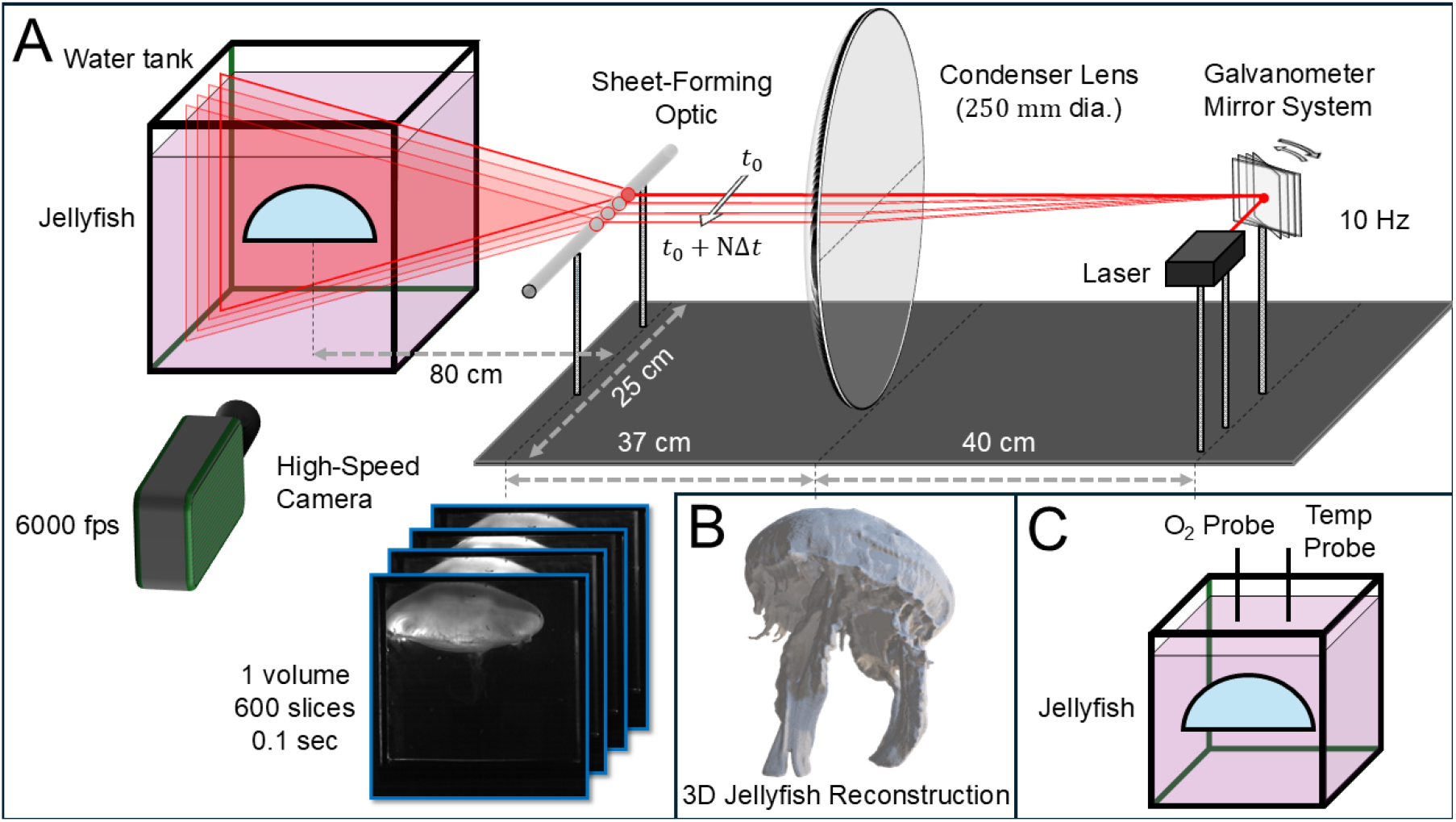
Laser scanning system for volumetric reconstructions. (A) Diagram of laser scanning system with laser, galvanometer rotating at 10 Hz, 250 mm diameter condenser lens to redirect the laser normal to the scanning direction, cylindrical glass rod to form the laser beam into a thin sheet, water tank, and jellyfish. As the laser scanned across the jellyfish, a high speed camera recorded an image stack, or “volume”, of 600 slices every 0.1 s. (B) A representative 3D volumetric reconstruction of a live jellyfish. (C) Closed system respirometry setup used to validate laser scanning technique. Jellyfish oxygen consumption was measured using an O_2_ probe and temperature probe over time in a sealed chamber. Figure adapted with permission from Fu et al., 2021.

### Validation of volumetric reconstruction

Experiments were performed to characterize the accuracy and precision of the laser scanning technique. To measure the relationship between volumetric reconstructions and wet weight of jellyfish, 10 jellyfish were scanned and weighed to compare wet weight (WW) and laser scanned weight (LSW). Each jellyfish was first scanned using the procedure described in this section. Then, they were carefully removed from the tank, patted dry to remove additional water, and weighed on an electric balance (Eosphorus Sf-400C, 600 g range, 0.01 g precision).

### Energetic consumption from volumetric changes

This laser scanning technique used repeated laser scans and volumetric reconstructions over several hours to convert a change in volume due to catabolysis to an average energy consumption rate for each jellyfish. In order to validate this conversion, simultaneous laser scanning and closed system respirometry experiments were performed. The jellyfish was scanned and reconstructed using the procedure described in this section and immediately transferred to a 15.1 liter closed respirometry chamber filled with artificial seawater at 35 ppt (Fig. 1C). An oxygen probe (OPTO-430, Unisense, Denmark) and temperature sensor (TEMP-UNIAMP-7, Unisense, Denmark) were used to continuously measure the oxygen concentration at a sampling frequency of 0.1 Hz. After 24 hours, the animal was removed, scanned, and the artificial seawater in the respirometry chamber was flushed and replaced. The animal was returned to the respirometry chamber for a second 24 hour respirometry measurement. Finally, at 48 hours from experiment initiation, the animal was removed, scanned, and the experiment was completed. The respirometry chamber was chosen to be large enough that the animal avoided hypoxic conditions over the course of the experiment. This experiment was repeated for 3 animals. Oxygen consumption in an empty respirometry chamber was measured and background oxygen consumption was subtracted from experimental results. A linear fit to the data was used as the average respiratory rate (RR). These values were compared with literature values for *Aurelia aurita* wet weight (WW) given by

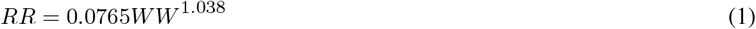

(Uye and Shimauchi 2005).

The method to convert the measured change in laser-reconstructed animal size to an energetic rate was based on earlier animal physiology work (Ikeda et al., 2000). The respiratory rate is defined as

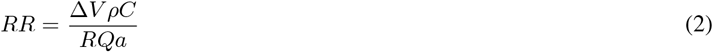

in terms of the change in volume of the animal Δ*V*, tissue density *ρ*, tissue carbon content *C*, respiratory quotient *RQ*, and a unit conversion term *a*. The change in volume Δ*V* was measured using the laser scanning method described in this section. Since jellyfish are neutrally buoyant and approximately 96% water (Uye and Shimauchi, 2005), *ρ* was assumed to be equal to the density of seawater *ρ* = 1024 kgm^-3^ at 35 PPT and 21°C. We experimentally measured the tissue carbon content C of 9 *Aurelia aurita* as described in the following section. The respiratory quotient *RQ* depends on the type of tissue metabolized and represents the ratio of carbon dioxide produced to oxygen consumed. For jellyfish, we used a range *RQ* = 0.8 *−* 0.9 since they metabolize primarily protein (Uye and Shimauchi, 2005; Ikeda et al., 2000). Uncertainty in these input parameters was propagated to provide error bounds for metabolic estimates. Previous work reports that the weight specific respiratory rate of *Aurelia aurita* does not change with starvation state (Frandsen and Riisgard, 1997) suggesting equation 2 does not need to be corrected for starvation state. We normalize all *RR* by animal *WW* in grams to allow comparisons across animal morphology.

### Carbon content

Tissue carbon content, *C*, was experimentally measured for 9 animals to measure body composition changes while undergoing catabolysis. The animals were transferred to a clean 108 liter artificial seawater tank without food. Three animals were randomly selected for removal at each timestep of 0 hours, 25 hours, and 50 hours after experiment start. Each animal was patted dry, *WW* measured on an electric balance, and transferred to an electric oven where they were dried at 60-65°C until a constant weight was achieved. Dried samples were pulverized in a mortar and pestle and weighed. Samples were then analyzed for carbon and nitrogen content on an elemental analyzer (Santa Barbara Marine Science Institute Analytical lab, CEC 440HA, Control equipment Corp, MA, USA). These experiments did not show statistically significant changes in carbon content over 50 hours without feeding. We found an average carbon content of 0.105% ± 0.026% of *WW*, which is not statistically different from the 0.13% ± 0.06% reported in the literature (Uye and Shimauchi, 2005). We used the value measured in the present experiments for the laser scanning method calculations. Analysis showed an average carbon to nitrogen ratio of 3.64 *±* 1.26 which is not statistically different from the literature value of 3.71*±* 1.63 (Uye and Shimauchi, 2005).

### Electrical stimulation for continuous swimming

Figure 2A shows the electronics for onboard electrical stimulation embedded in a model jellyfish. The system consists of a swim controller enclosed in a custom 5.10 ± 0.03 cm diameter electronics housing, two electrodes embedded in the jellyfish muscle tissue, a threaded plastic rod, and a ballasting cap. The swim controller includes a small processor (TinyLily, TinyCircuits, Akron, OH, USA) and a 250 mAH lithium-polymer battery cell (GM602025-PCB, PowerStream Technology Inc. Orem, UT, USA). All parts of the electronics housing and ballast were fabricated via stereolithography (Form 3 Color Base Resin) on a Form 3+ resin printer (Formlabs, Somerville, Massachusetts). The swim controller housing utilized a threaded cap with two O-rings for waterproofing at high pressure. Wire coated in perfluoroalkoxy was threaded through small holes in the housing and connected to platinum rods (A-M Systems, Sequim, WA, USA) embedded in the animal muscle tissue. The pass-throughs were sealed with epoxy and an LED was connected in series to each electrode to provide visual confirmation of stimulation. The ballasting cap passively maintains vertical downward swimming and could house future sensors. See previous work (Anuszczyk and Dabiri, 2024) for more details on electronics and ballasting cap design and impact on swimming performance. Figure 2B shows the electronics stimulating a swimming jellyfish in the lab environment. The simultaneous laser scanning and respirometry experiments described in this section were additionally performed with 3 electrically stimulated animals.

**Fig. 2.**
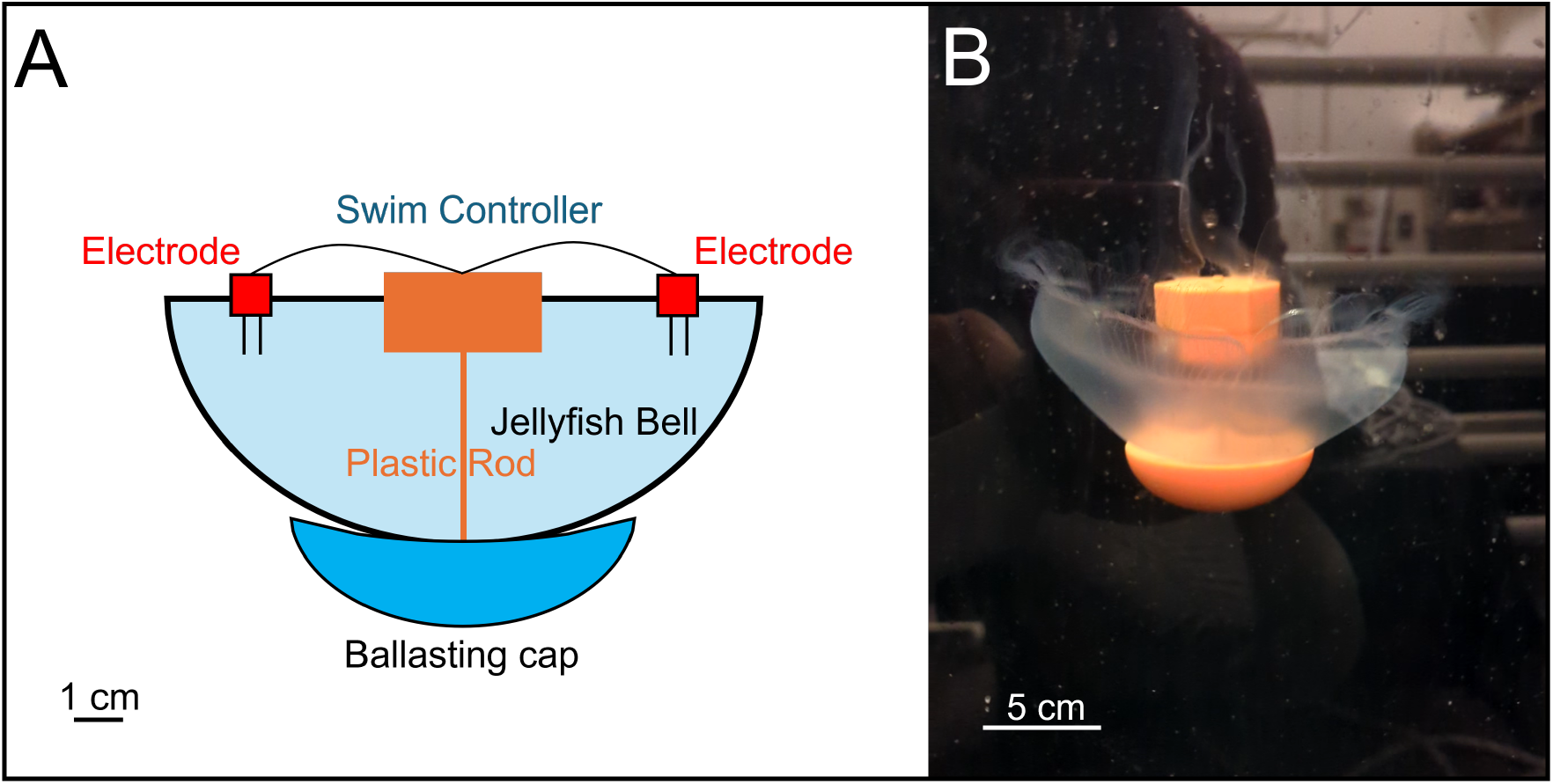
Jellyfish with embedded electronics for electrical stimulation at constant pulse rate. (A) Diagram of electronics with swim controller housing TinyLily microprocessor with 250 mAH battery, two electrodes embedded in bell margin stimulating at 0.5 Hz, plastic rod to connect components, ballasting cap to passively maintain downward vertical swimming, and 1 cm scale bar. (B) Electronics embedded in live jellyfish during continuous swimming experiment with 5 cm scale bar. This figure was adapted from Anuszczyk and Dabiri, 2024.

### Long duration stimulated free-swimming experiments

The laser scanning method was utilized to measure the free-swimming energy consumption of electrically stimulated, continuous swimming jellyfish in a 6-m tall, 13,600-liter vertical tank (Figure 3A). The tank was filled with artificial seawater balanced at 35 PPT and 21°C and has a continuous filtration pump which was turned off at least 2 hours before these experiments. This tank is divided into a test section on the right, where experiments are conducted, and a recirculation region on the left with water pumps (A1200, Abyzz, Germany). This enabled continuous flow, shown with red arrows, through the test section functioning as a “flow treadmill”. During long-duration swimming experiments, a jellyfish was equipped with stimulation electronics, weighted to maintain slight positive buoyancy with a stable downward orientation, and released at the top of the tank to swim downwards. A camera microprocessor (RT1062 Cam, OpenMV, Atlanta, GA, USA), set back 0.86 m from the tank, was programmed with a custom MicroPython computer vision algorithm. This computer vision camera tracked the animal location during continuous swimming and was programmed to interface with the water pumps while running a Proportional-Integral-Derivative control loop. Figure 3B shows the view from the camera of the middle section of the tank as a jellyfish was swimming downwards. As the animal descends in the tank, the controller continuously adjusts the flow opposing the jellyfish with an optimization goal of keeping the animal within the middle section of the tank and in view of the camera. The computer vision camera was programmed to track both the animal location and the pump power. Particle Image Velocimetry (PIV) was conducted to calibrate pump power to pump flow speed and confirm flow uniformity. Animal swimming speed was found by combining the movement of the animal in the lab frame with the pump flow speed. During the course of the 50 hour experiments, the animal was occasionally out of frame for brief periods. These periods were removed when calculating swimming metrics for speed and distance.

**Fig. 3.**
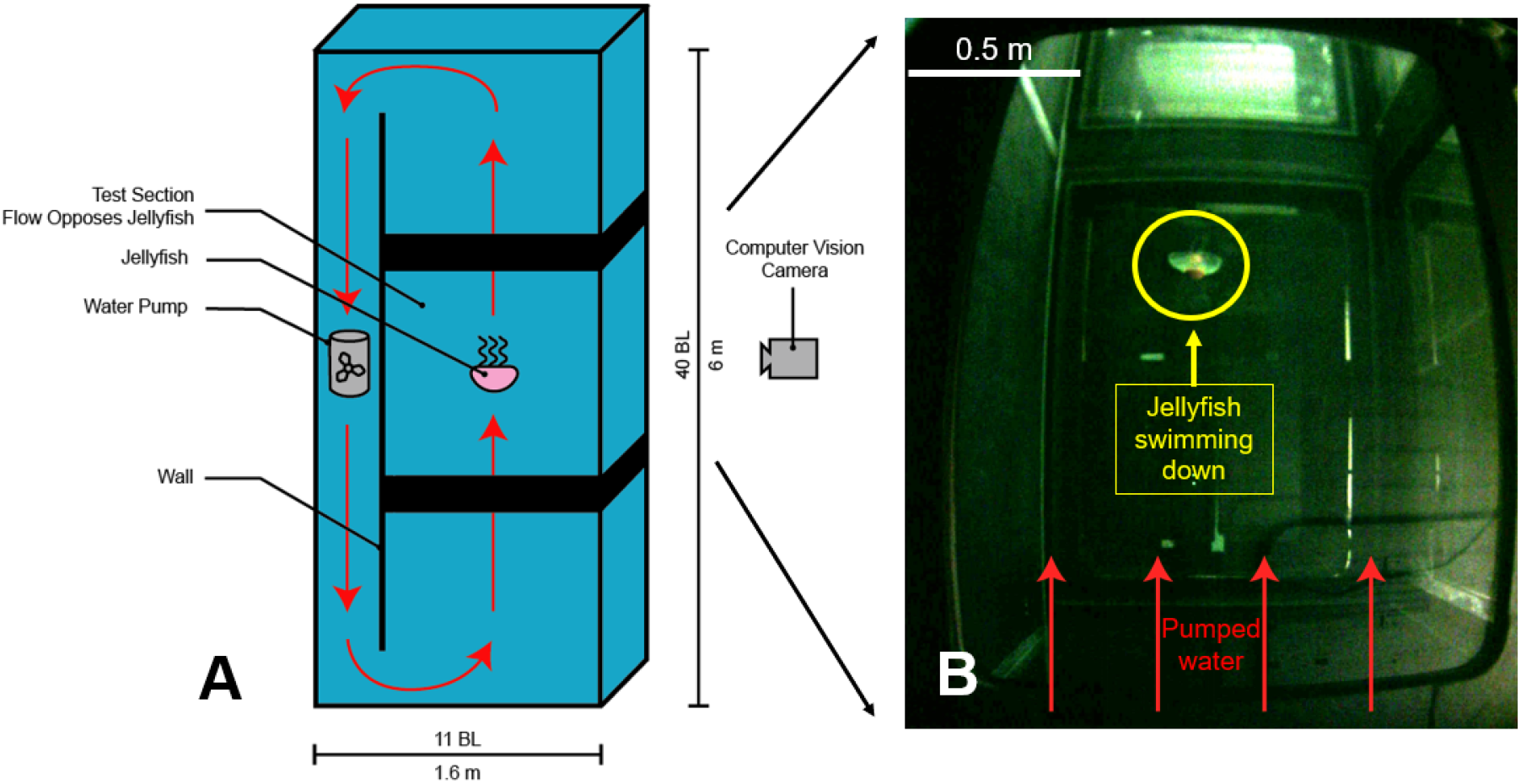
Free-swimming tank for long-duration experiments. (A) Diagram of 6 m by 1.6 m artificial seawater vertical tank facility. Water pumps in the recirculation area on the left generated flow (shown with red arrows) which opposed free-swimming jellyfish. A computer vision camera tracked the swimming animal. (B) The view from the computer vision camera of the middle section of the tank as a jellyfish, circled in yellow, swam downward. As the jellyfish descends, the camera ran a PID control loop interfacing with the water pumps to control the flow keeping the animal in view of the camera.

### Basal metabolism

The total metabolism of free-swimming animals comprises both metabolic requirements to maintain swimming and basal metabolism for other physiological processes. Additional experiments were conducted to quantify the basal metabolic component of the total metabolic measurements. To the best of our knowledge, these are the first measurements of the basal metabolism of jellyfish without animal movement. Basal metabolic experiments were performed using a 2.55 liter glass respirometry chamber and the oxygen probe and temperature sensor described in this previous section with 6 animals. Two methods were investigated successively: anesthetization with magnesium chloride (MgCl_2_) to suppress muscle movement and excision of the nerve centers, or rhopalia, of the animal in order to arrest endogenous swimming signals. Excising rhopalia has been shown to arrest normal contractions (Romanes 1885; Satterlie, 2002) and the animals were observed to not contract for the duration of this experiment after excision. Each animal was transferred to the respirometry chamber filled with artificial seawater where they underwent five distinct treatments of 1 hour each. This enabled testing of both methods and comparison with natural respiration rates. First, the chamber was closed and normal respiration rate was measured. Next, the animal was removed and MgCl_2_ (AC413410025, Fisher Scientific, Waltham, MA, USA) was added at a molarity of 0.3 M. The MgCl_2_ was mixed for 5 minutes until dissolution, and the animal was added for the anesthetization test. Upon completion, the water was then exchanged for clean artificial seawater and a second normal respiration rate was measured to ensure the animal had fully recovered from anesthetization. The same animals were then used to test the excision method. Respiratory rate was measured again for the immobilized jellyfish after excision. The animal was again removed from the tank, the water was replaced with the MgCl_2_ solution, and respiratory rate was measured combining both anesthetization and excision methods. While both anesthetization and excision methods were tested, we report results from the excision method here due to potential confounding impacts of MgCl_2_ with animal tissue on oxygen consumption.

### Swimming metabolism model

We derived a physics-based quasi-steady swimming metabolism model based on previous work (Dabiri et al., 2006; Acuña et al., 2011). Here, we model the jellyfish energy consumption 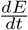 as the sum of the basal metabolism *Rb*, the power expended setting the surrounding water into motion while swimming *P*_*wake*_, and the hydrodynamic drag on the free-swimming animal *P*_*drag*_

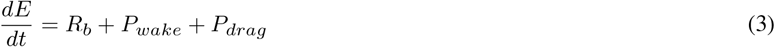

We modele the water motion created by the animals while swimming as primarily comprising a vortex ring. Following previous work (Dabiri et al., 2006), we approximate its kinetic energy as that of an equivalent oblate spheroid of water set into motion each time the jellyfish pulses and generates a vortex ring. We use the classical kinetic energy formula for the kinetic energy contained in the wake,

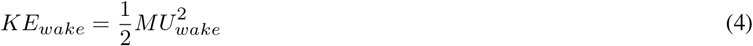

where *M* is the mass of water set in motion and *U*_*wake*_ is the speed at which this wake convects. The mass is given by *M* = *ρV* where seawater density *ρ* = 1024 kgm^-3^ at 35 PPT and 21°C, and the volume 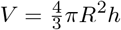 where R is the radius of the animal and h is the semi-minor axis of the spheroid. As the vortex ring is accelerated, it accelerates the surrounding fluid necessitating an added mass term *C*_*am*_. We also introduce a propulsive efficiency term *η* to account for energy lost in physiological processes that convert body motion to fluid motion leading to

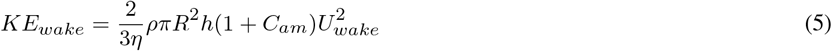

Power is the time rate of change of kinetic energy. *Aurelia aurita* generate two vortex rings with each swimming cycle (Gemmell et al., 2013). Thus, we assume that the kinetic energy for two vortex rings is expended with each swimming cycle. Hence,

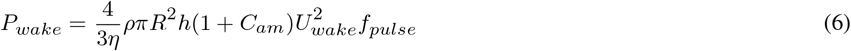

where *f*_*pulse*_ is the pulse frequency. We model the drag on the free-swimming jellyfish as the product of the drag force and the swimming speed divided by propulsive efficiency

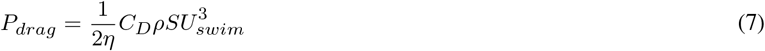

with *C*_*D*_ the drag coefficient for the jellyfish, *S* the projected planform area in the direction of swimming, and *U*_*swim*_ the swimming speed of the animal. Thus, combining equations 8 and 6 into 3 becomes

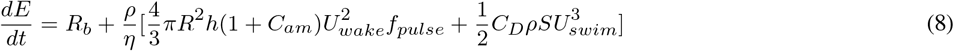

Jellyfish geometry was measured to determine *R*, and *h* was estimated as *h* = 0.4*R*. Stimulated pulse rate was set by the electronics at *f*_*pulse,stim*_ = 0.5 Hz, and previous work found jellyfish natural pulse rates of *f*_*pulse,nat*_ = 0.16 Hz (Xu and Dabiri, 2020). We estimated a range of *η* = 0.2 *−* 0.4 from previous work (Acuña et al., 2011). *C*_*am*_ was assumed to be 1, and for a flat plate *C*_*D*_ = 1.12 (Hoerner 1965). The wake convection speed *U*_*wake*_ was approximated as the contraction velocity of the bell margin, measured by analyzing a naturally pulsing animal and *U*_*swim*_ for the free-swimming experiments was measured as described in this section. To compare with respirometry data, we assumed 447 kilojoules per mole of oxygen consumed for protein, 12/22.4 is the weight of carbon in 1 mole of carbon dioxide (Ikeda et al., 2000), and there are 2.4 *×* 10^4^ ml per mole of oxygen (Garcia and Gordon, 1992). To normalize by wet weight mass, the relation

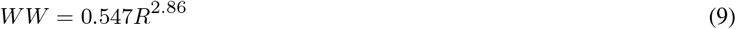

was used (Uye and Shimauchi, 2005).

## RESULTS

### Laser scanning technique validation

We compared wet weight (*WW*) to laser scanned weight (*LSW*) to validate this technique. Figure 4A shows the linear relationship between the measured *WW* and *LSW* of *WW* = 0.63*LSW −* 17.66 with *R*^2^ = 0.9256 for 10 animals of different sizes. The *LSW* overestimated the *WW* of the animal, likely due to the scattering of the laser sheet through the tissue. This well-defined relationship enabled us to convert accurately between *WW* and *LSW*.

**Fig. 4.**
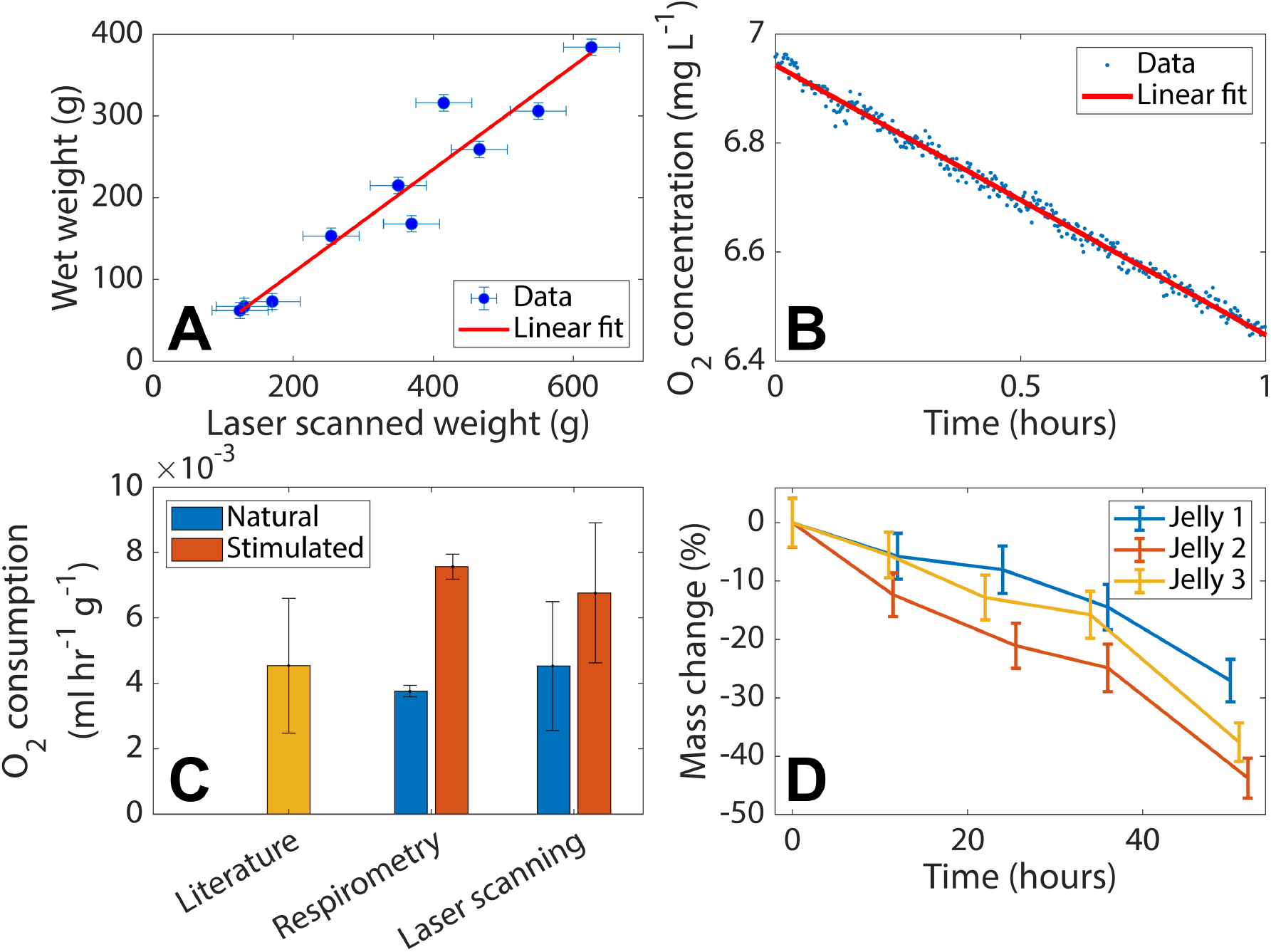
Validation of laser scanning technique for energy measurements. (A) Linear relationship between wet weight (WW) and laser scanned weight (LSW) of *WW* = 0.63*LSW* − 17.66 for 10 animals. Error bars represent 1 standard deviation for each axis. (B) A sample 1 hour closed system respirometry experiment with linear fit and R-squared value of *R*^2^ = 0.9921. This high R-squared value confirms the linearity of jellyfish oxygen consumption over 1 hour. (C) Literature values for natural animals compared with simultaneous closed system respirometry and laser scanning. There is no statistical difference between the oxygen consumption rates found with the two different techniques. Oxygen consumption given in terms of ml O_2_ per hour per g wet weight. (D) We deployed this technique to measure free-swimming metabolism during 50 hour experiments for 3 jellyfish. The jellyfish lost 36% ± 8.4% of their mass on average over the course of this experiment.

Simultaneous laser scanning and respirometry experiments confirmed the accuracy of this method. Figure 4B shows a representative 1 hour respirometry experiment. A linear fit to this data has an R-squared value of *R*^2^ = 0.9921 and illustrates the linear trend in oxygen consumption over the course of these experiments. The literature bar represents the wet weight normalized oxygen consumption for an animal of the same size as those used in the respirometry and laser scanning experiments shown in Figure 4C. This was found using equation 1 with the error given as ± 1 standard deviation. The respirometry bar compares the animal’s natural and electrically stimulated respiratory rate we measured in the respirometry chamber with the error given by ± 1 standard deviation of the linear fit to the respirometry data. The laser scanning bar in Figure 4C also compares naturally swimming animals to electrically stimulated animals measured with the laser scanning method. This error represents ± 1 standard deviation and combines the errors in scanning and equation 3. Electrically stimulated animals were found on average to consume 73% ± 32% more energy than naturally swimming animals in this confined configuration.

These experiments validate that the observed laser scanning respiratory rate from catabolysis, calculated using equation 2, correspond with measurements from the respirometry chamber. The blue “Natural” bars have no statistically significant difference with a two-tailed t-test p-value of p=0.44. Similarly, the red “Stimulated” bars have no statistically significant difference with a two-tailed t-test p-value of p=0.60. This confirms that although the laser scanning method has a higher error compared to the respirometry chamber, the two methods are in agreement and the technique is suitable for further experiments.

### Long duration stimulated free-swimming experiments

We utilized this laser scanning technique to measure the free-swimming metabolism of 3 electrically stimulated continuously swimming jellyfish in a 6-m tall vertical tank. Figure 4D shows the mass change of each animal as a percent of the mass at the previous timestep. The error here is found by scanning an animal multiple times and taking the mean and standard deviation of the resulting measurement. The continuously swimming animals lost a mean of 36% ± 8.4% of their mass over the course of 50 hours which is in line with previous results on starvation (Ishii and Bamstedt, 1998). The starting masses of the animals found from scanning were *m*_*jelly*,1_ = 470 *±* 19 g, *m*_*jelly*,2_ = 347 *±* 14 g, and *m*_*jelly*,3_ = 536 *±* 21 g. We found that the animals swam a mean distance of 2.55 ± 0.46 km while in view of the computer vision camera at speeds of 2.44 ± 0.60 cm s^*−*1^. Previous results showed similar electrically stimulated swimming speeds over much smaller timescales on the order of minutes (Anuszczyk and Dabiri, 2024).

Free-swimming experiments showed increased energy consumption. Figure 5 compares the energetic costs of swimming jellyfish in the three different configurations tested. The blue bars represent the total metabolism. In the “Natural enclosed” configuration and the “Stimulated enclosed” configuration, this was measured in the respirometry chamber with the oxygen probe, and in the “Stimulated free-swimming” configuration using the laser scanning technique in the vertical tank. We found that the free-swimming animal consumes 2.5 times more energy, normalized by body mass, than the stimulated animal in the enclosed respirometry chamber. This increase in energy consumption is likely due to a combination of hydrodynamic effects. Because hydrodynamic power scales as the cube of swimming speed (Daniel 1983), small increases in speed result in disproportionately higher energetic costs. The increase in swimming speed from nearly zero in the enclosed configurations to 2.44 ± 0.60 cm s^*−*1^ in the free-swimming configuration would thus increase the hydrodynamic power expended. In addition, recirculation in a small respirometry chamber reduces the force required to contract the bell, suggesting that the same movement might require more energy during free-swimming.

**Fig. 5.**
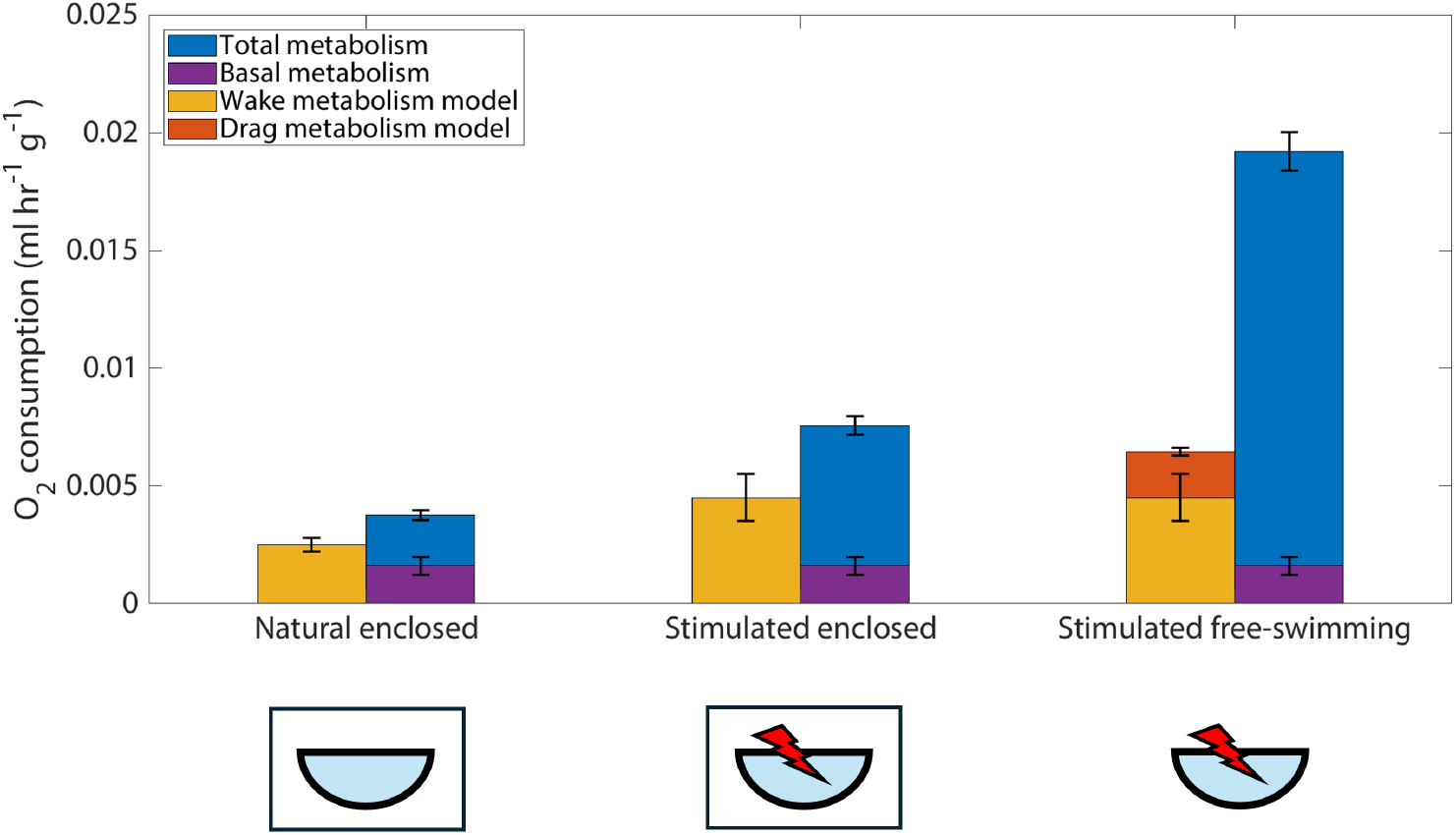
Free-swimming energy consumption and metabolism model. Total metabolism, shown in blue, in enclosed configurations here was measured using closed respirometry, free-swimming total metabolism was measured using the laser scanning technique. Stimulated free-swimming jellyfish consume 2.5 times as much energy as stimulated enclosed animals. Diagrams represent jellyfish in each configuration. We measured basal metabolism, shown in purple, by excising rhopalia as 42% ± 10% of jellyfish natural swimming energy consumption. The wake metabolism model, in yellow, representing the energy to set water in the wake into motion, underestimates energy consumption. Adding a steady drag metabolism term, in red, for the free-swimming jellyfish continues to underestimate the energy consumption. Oxygen consumption given in terms of ml O_2_ per hour per g wet weight.

**Fig. 6.**
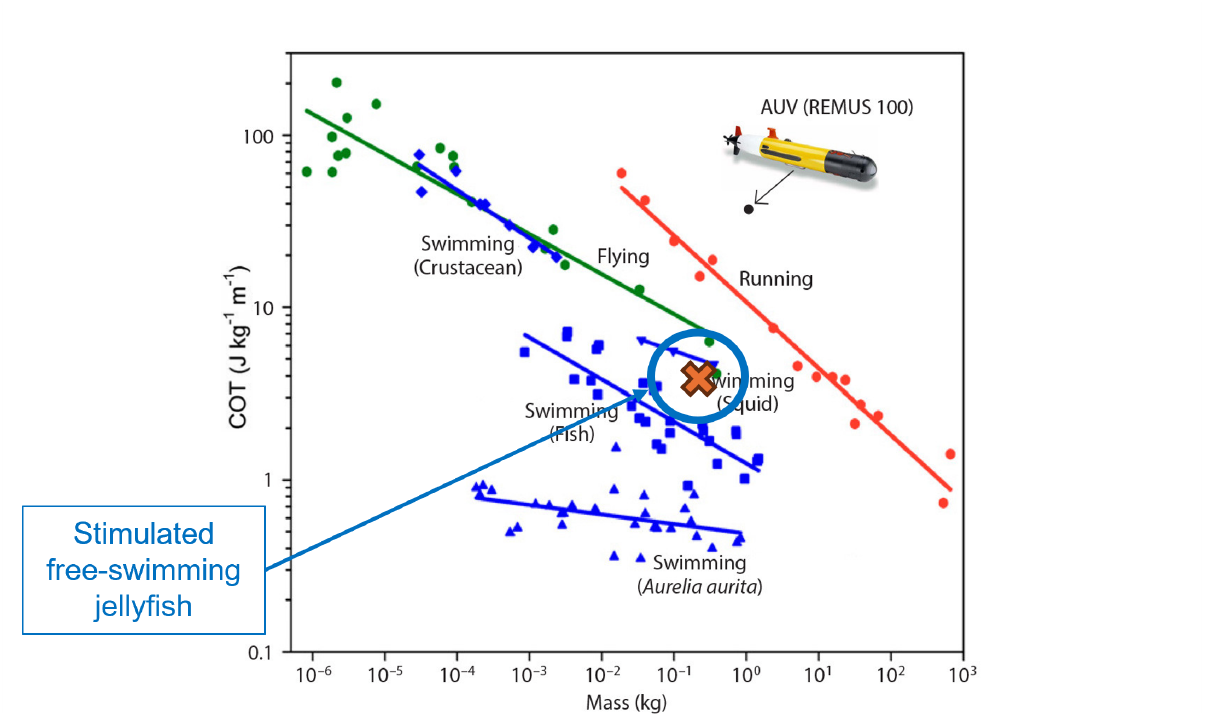
Cost of Transport of stimulated jellyfish compared to other animals. While natural jellyfish are the most efficient metazoans (Gemmell et al., 2013), the increase in energy consumption found with the laser scanning technique increases the COT. Thus, these jellyfish have a COT comparable to swimming fish, as shown with the orange x. This COT is more than an order of magnitude smaller than a representative autonomous underwater vehicle (AUV), the REMUS 100. Figure adapted with permission from Gemmell et al., 2013.

We found an average basal metabolism of *R*_*b*_ = (1.6 *±* 0.4) *×* 10_*−*3_ ml O_2_ hr^-1^ g^-1^ which constitutes 42% ± 10% of jellyfish natural swimming energy consumption based on our experiments with rhopalia excision. This is shown in figure 5 in purple for each test configuration. While there is little data on basal metabolism of gelatinous zooplankton, this result is comparable to some fish, e.g. tuna species *Katsuwonus pelamis*, with a basal metabolism that constitutes 58% of natural swimming total metabolism (Brill 1987). Jellyfish are known to be among the most efficient metazoans in terms of cost of transport (COT) defined as the mass-specific energy per distance traveled (Gemmell et al., 2013). Their low basal metabolism compared to other species suggests that their efficiency is not only due to efficient swimming as previously proposed. Instead, the low COT is also due to highly efficient essential physiological functions that make up basal metabolism such as digestion and reproduction.

Our swimming metabolism model is shown in Figure 5 in orange alongside each configuration with error bars representing a range of propulsive efficiencies. The drag term is only shown for the free-swimming animal in red as described in Methods. This model generally captures the trends of the “Natural enclosed” and “Stimulated enclosed” configurations. However, it struggles to account for the large increase in energy consumption of the free-swimming animal necessitating additional investigation.

## DISCUSSION

We have presented measurements of electrically stimulated free-swimming jellyfish total metabolism that we found to be 2.5 times higher than similarly stimulated animals in a closed respirometry chamber. This method of measuring metabolic rates utilizes laser scanning of jellyfish which we calibrated and validated through additional experiments. Basal energy consumption measurements showed that jellyfish consume 42% ± 10% of total energy consumption for physiological functions other than swimming.

Converting the increased energy consumption into cost of transport (COT) shows that free-swimming stimulated jellyfish energy consumption is on a similar scale as swimming fish (Figure 7). Since this study utilized increased pulse rates, this COT suggests that one contributor to natural jellyfish efficiency is less frequent pulse rates which often correspond to periods of coasting between pulses. Further investigation by conducting metabolic experiments with lower frequency stimulation could clarify the impact of pulse rate on COT.

Experimentally measuring total metabolism of electrically stimulated jellyfish enables further work developing this platform for ocean exploration. Previous work found portable electrical stimulation combined with an exumbrellar ballasting cap increases animal swimming speeds by up to 4.5 times natural swimming speeds (Xu and Dabiri, 2020a; Anuszczyk and Dabiri, 2024). These stimulated animals are capable of carrying scientific payloads of up to 105% of animal mass in this ballasting cap, enabling the carry of a variety of scientific sensors to study the biogeochemistry of the ocean (Anuszczyk and Dabiri, 2024). Field tests have demonstrated vertical profiling capability in oceanic conditions near the top of the water column (Xu and Dabiri, 2020b; Rutledge et al., in prep). Our results show that stimulated animals can swim more than 2.55 km or 15,000 body lengths which could enable deeper deployment as ocean sensors. Since these continuous swimming experiments were performed in the top 6-m of the water column, deeper deployments would be contingent upon experiments to ensure both the animal and electronics can sustain higher pressures. While the increased COT is higher than natural animals, this COT is more than an order of magnitude smaller than a representative AUV, and thus far more efficient than traditional methods of ocean exploration.

One remaining question on the feasibility of deep ocean deployments of this platform is if zooplankton distribution and animal clearing rates will provide sufficient feeding opportunities to avoid significant catabolysis. Zooplankton distribution research in the Beaufort sea of the Arctic Ocean suggests that biovolume peaks at around 50-100 meters depth. However, some zooplankton such as *Chaetognaths* and other gelatinous zooplankton were found at high biovolume densities at 1000 m, the depth limit of this study (Forest et al., 2012). This suggests that jellyfish deployed to these depths may find different but sufficient feeding opportunities. Future work could incorporate feeding at different spatial densities into similar studies to experimentally study jellyfish ability to feed during electrical stimulation.

We developed a model that generally captures the energy consumption trends observed experimentally. After subtracting basal metabolism, the measurements of energy consumption increase by a factor of 2.8 between the natural and stimulated enclosed configurations. This closely corresponds with the increase in pulse rate 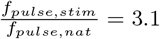 suggesting that energy consumption per pulse is relatively constant across pulse rates in an enclosed configuration. Therefore, this model underestimated the energetic consumption associated with each pulse, in all configurations, potentially due to the induced unsteady flow field. While the majority of the vorticity is concentrated in vortex rings, other wake energy is distributed spatially highlighting the complicated nature of jellyfish wake flow dynamics (Gemmell et al., 2015). Additional work could refine this model by taking into account more detailed wake reconstructions and vortex measurements. Conducting high-resolution PIV on the wake of natural and stimulated organisms could contribute to more accurate models. Muscle inefficiency during stimulation could also lead to increased energy consumption. Previous work, using a simple drag model, has explained low energy consumption model predictions by assuming a higher basal metabolism than that measured in this study (Acuña et al., 2011).

Although this study focused on *Aurelia aurita*, this method holds potential for application to other marine species. Recent work has achieved detailed laser scans of *Nanomia bijuga* and *Cystisoma* (Daniels et al., 2023). Our work extends these findings with automated reconstructions of larger transparent animals. Scattering and attenuation within jellyfish tissue limit laser transmission, obscuring features on the far side of the volumetric reconstruction. Reconstructions could be utilized for comparative studies of animal morphology, biomechanics, and behaviour given sufficient camera frame rates to limit movement between successive laser scans. This metabolism technique could be extended to a broad range of gelatinous zooplankton but would require modification for application to more complex organisms. While jellyfish have little tissue type variation, application to some vertebrates, e.g. bony fishes, would require tracking tissue type lost through catabolysis to account for differing chemistry and carbon content.

## Acknowledgements

The authors would like to thank Cabrillo Marine Aquarium and Aquarium of the Pacific for providing jellyfish used in these experiments. The authors also acknowledge helpful conversations with Lea Goentoro, Michael Dickinson, Morteza Gharib, John Costello, Sean Colin, Brad Gemmell, Emily Carrington, and Mark Denny.

## Contribution

S.R.A. and J.O.D. conceived of project, S.R.A. conducted experiments, S.R.A. and J.O.D. analyzed results and wrote paper.

## Data availability

Raw image data available upon request.

